# Development of a drug repressible β-catenin mutant mouse to examine effect of activated β-catenin signaling on long-term maintenance of mammary tumor growth

**DOI:** 10.1101/588459

**Authors:** Jennifer L. Gorman, Jessica R. Adams, Emma M. Jones, Sean E. Egan, James R. Woodgett

## Abstract

Stabilized β-catenin expression is a well described initiator of mammary tumorigenesis in the mouse and elevated nuclear expression of this protein has been observed in human triple-negative tumor samples. However, the importance of stabilized β-catenin to continued tumor growth after initiation, and in the context of other driver mutations, has yet to be elucidated. To ascertain the importance of stabilized β-catenin after tumor initiation, we generated a novel transgenic mouse model, utilizing the tet-off system, to control expression of a stabilized mutant of β-catenin. Pups from early litters carrying one allele of regulatable stabilized β-catenin at the Rosa26 locus, but not targeted by Cre, were smaller in size compared with wildtype littermates and also developed skin lesions, requiring euthanasia. Maintenance of breeding cages on doxycycline chow allowed for healthy transgenic pups to survive past 3 weeks of age and 6 animal cohorts were established. We used two different mammary Cre strains, WAP-Cre and MMTV-NLST-Cre, crossed with regulatable, stabilized β-catenin (bcatFL) animals, as well as animals expressing mutated p53-R270H. While mammary tumors developed in animals expressing Cre-targeted regulatable bcatFL, the penetrance and growth kinetics were lower than observed with other mammary mouse models expressing stabilized β-catenin. Since fewer animals developed tumors, we focused analysis on the effect of turning off stabilized β-catenin expression following tumor initiation. In the animals examined, a delay in tumor growth, but not regression, was observed in animals expressing Cre-targeted bcatFL as well as those expressing both bcatFL and p53-R270H. These results indicate that in a less aggressive β-catenin model, additional mutations likely provide independence from the initiating event.

## Introduction

Wnt pathway activation induces stabilization of cytoplasmic and then nuclear translocation of β-catenin, a process which drives cell proliferation, while also promoting cell self-renewal and inhibiting cell differentiation (1, 6, 18). The role of activated β-catenin signaling in the initiation of mammary tumorigenesis in the mouse has been well established. Whether through N-terminal deletion (ΔN57, ΔN89 or ΔN90) or Cre-mediated exon 3 removal, activation of β-catenin leads to tumor formation, with latency and penetrance dependent upon which mammary cell populations are targeted (11, 13-15, 22, 23). In mammary luminal cells targeted by MMTV driven expression, mammary gland hyperplasia is observed with the development of adenocarcinomas between 7-12 months of age (11, 13). If the Whey Acidic Protein (WAP) promotor is used to target expression, squamous metaplasias develop after glandular transdifferentiation (14, 15). Finally, targeted expression to basal cells leads to the development of aggressive carcinomas, but only in multiparous animals. Nulliparous females show gland transdifferentiation but no tumor development (23). While metastasis is not documented in mice solely expressing activated β-catenin, crossing Δ90-β-catenin mice with p53 +/− mice leads to the development of lung metastasis in just over a third of the animals which developed mammary tumors (19). Loss of an allele of p53 also led to increased tumor burden and decreased latency of mammary tumor formation (19). Wnt signaling is also associated with breast tumor formation, as human breast biopsies show elevated nuclear levels of β-catenin, especially in the highly invasive “triple negative” subtype (9, 23).

While the role of activated β-catenin in mammary tumor initiation is well documented, it’s role in the long-term maintenance of tumor growth has yet to be ascertained. Studies on other drivers of tumorigenesis in mice have revealed differing roles once tumor initiation has occurred. Ras-driven melanoma tumor formation could be reversed upon H-Ras-V12D withdrawal (4) and K-Ras driven lung tumors also regressed upon inactivation of the oncogene, with or without the presence of secondary mutations (p53 or Ink4A/Arf) (8). c-myc induced acute myeloid leukemia regresses upon loss of c-myc overexpression, resulting in normal hematopoiesis (7), as well, osteosarcoma driven by myc not only regressed following inactivation, but subsequent re-expression of myc did not result in re-initiation of tumor growth (12). While removal of the driving oncogene in other types of tumors resulted in long-term tumor regression, in mammary tumorigenesis this regression was not maintained in some models. Removal of activated HER2/Neu following tumor establishment was shown to induce tumor regression in both xenografted NIH3T3-her2 expressing cells and spontaneous mammary tumors arising in HER2/Neu mice (16, 20). However, continued suppression of Neu expression did not prevent the regrowth of tumors, which had no evidence of elevated Neu expression (16, 20). This was also observed with Wnt-1 driven mammary tumors, which regressed upon Wnt1 removal, but eventually Wnt1-independent tumors emerged (10). In c-myc driven mammary tumors, driver withdrawal lead to regression only in a subset of tumors that did not harbour a Ras family mutation (5). In tumors that did regress, recurrence was observed that was both myc-driven and myc-independent (3).

Given the importance of some oncogenic drivers to the long-term maintenance of tumor growth, we sought out to establish the importance of persistent, activated β-catenin to this process both with, and without, a secondary driving mutation. Here we report our generation of a regulatable mouse model of activated β-catenin utilizing the tetracycline machinery, in which doxycycline administration turns off transgene expression.

## Results

### Generation of regulatable stabilized β-catenin mouse model

Our regulatable stabilized β-catenin mouse model (bcatFL) required several features: the ability to tissue restrict expression, co-expression of stabilized β-catenin and luciferase to monitor tumor growth and metastasis, and the ability to turn expression on and off in the tissue of interest. To accomplish this, we first used a stabilized β-catenin cDNA which had previously been generated in our group through site-directed mutagenesis of 4 key serine/threonine residues (S33, S37, T41 and S45) to alanine to produce β-catenin that is resistant to phosphorylation-directed degradation. This stabilized β-catenin cDNA was then linked to a firefly luciferase gene via a T2a sequence. These sequences were cloned into the pTet-BigT vector downstream of a lox-stop-lox sequence, allowing for tissue restricted expression. This plasmid also contained the tetracycline machinery including the tetracycline transactivator (tTa), and a mini CMV/tetO promoter. Finally, this sequence was cloned into the ROSA26 PAM1 plasmid, so the finalized vector could be targeted to the endogenous Rosa26 locus. The Rosa26 promoter drives tTa expression, which binds to the tetO/mini-CMV promoter and, upon removal of the lox-stop-lox sequence, drives expression of firefly luciferase and stabilized β-catenin. When mice are fed doxycycline, tTa is blocked from binding to the tetO/mini-CMV promoter, thereby turning off transgene expression (Figure 1A)

**Figure 1.**
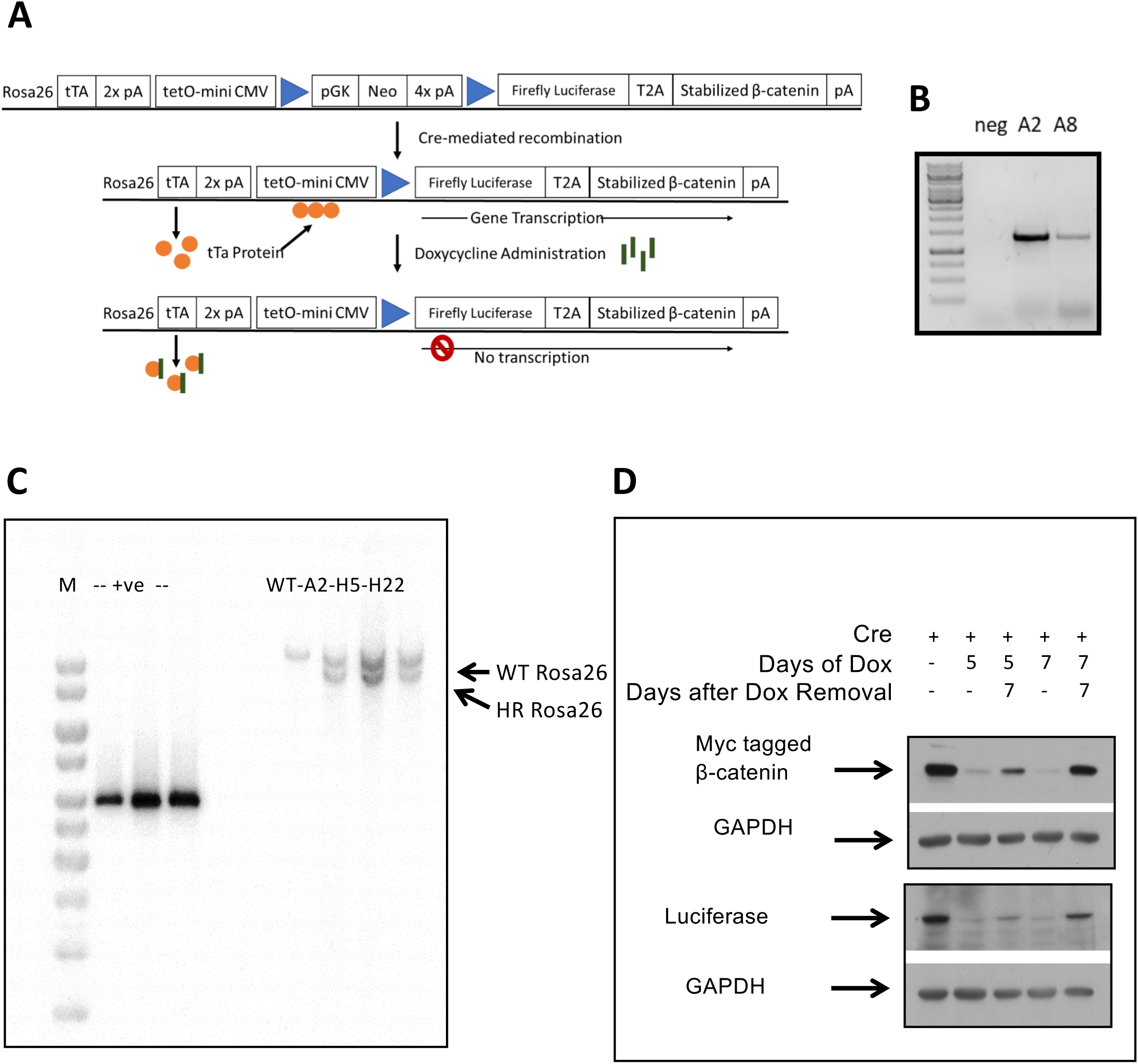
Generation of stabilized β-catenin-luciferase mouse model. A: schematic of the novel regulatable stabilized β-catenin-luciferase expressing mouse model: tTa, tetracycline transactivator; tet-O, tetracycline operator; CM, cytomegalovirus; Neo, neomycin resistance gene; pA, polyA. B: PCR screening of ES cell clones for proper insertion of bcatFL transgene at Rosa26 locus. C: Southern blot of ES cell clones positive for homologous recombination at the Rosa26 locus. D: immunoblot of ES cells demonstrating expression and regulation of myc-tagged β-catenin and luciferase expression following Cre transfection, with and without doxycycline administration.

The linearized targeting vector was electroporated into R1 embryonic stem cells and a total of 6 clones showed homologous recombination at the Rosa26 locus by PCR screening (Figure 1B), which was further confirmed in the A2, H5 and H22 clones by Southern blotting (Figure 1C). These clones underwent testing to prove that the system would work as expected. First, the clones were transfected with the pCAGGS-Cre plasmid to remove the lox-stop-lox sequence and tested for transgene expression. Strong expression of both luciferase and stabilized β-catenin (observed with blotting for the myc tag and β-catenin) was demonstrated for both the H5 and H22 clones (Figure 1D). The ability to turn off β-catenin expression was also tested. Incubation of cells with doxycycline for 5 days reduced both luciferase and stabilized β-catenin expression, and by 7 days little to no expression remained. Cells were also allowed to recover for 5-7 days after the removal of doxycycline and by day 5 faint expression of both transgenes was observed, with stronger expression 7 days after the cessation of doxycycline treatment. The H5 and H22 clones were then selected for diploid aggregation with ICR embryos. Following chimera breeding to both C57Bl/6 and FVB mice, germ-line transmission was observed from both clones.

### Development of skin condition in transgene positive pups

Pups displaying germline transmission were first screened for the presence of the β-catenin transgene via PCR (Figure 2A). However, from the first litters (on either C57Bl/6 or FVB) it was obvious which pups carried the transgenes. These pups were smaller than their wildtype littermates, and developed skin lesions on either their tail or in the peri-anal region (Figure 2B). The condition became so severe that mice were routinely sacrificed according to humane endpoint guidelines by 4 weeks of age. Tissue was collected from these mice to determine whether there was aberrant expression of the transgenes, even though the lox-stop-lox had not been removed. Western blotting of liver samples showed no expression of the transgenes (Figure 2C), and no luciferase expression was detected during bioluminescent imaging of these animals (data not shown).

**Figure 2.**
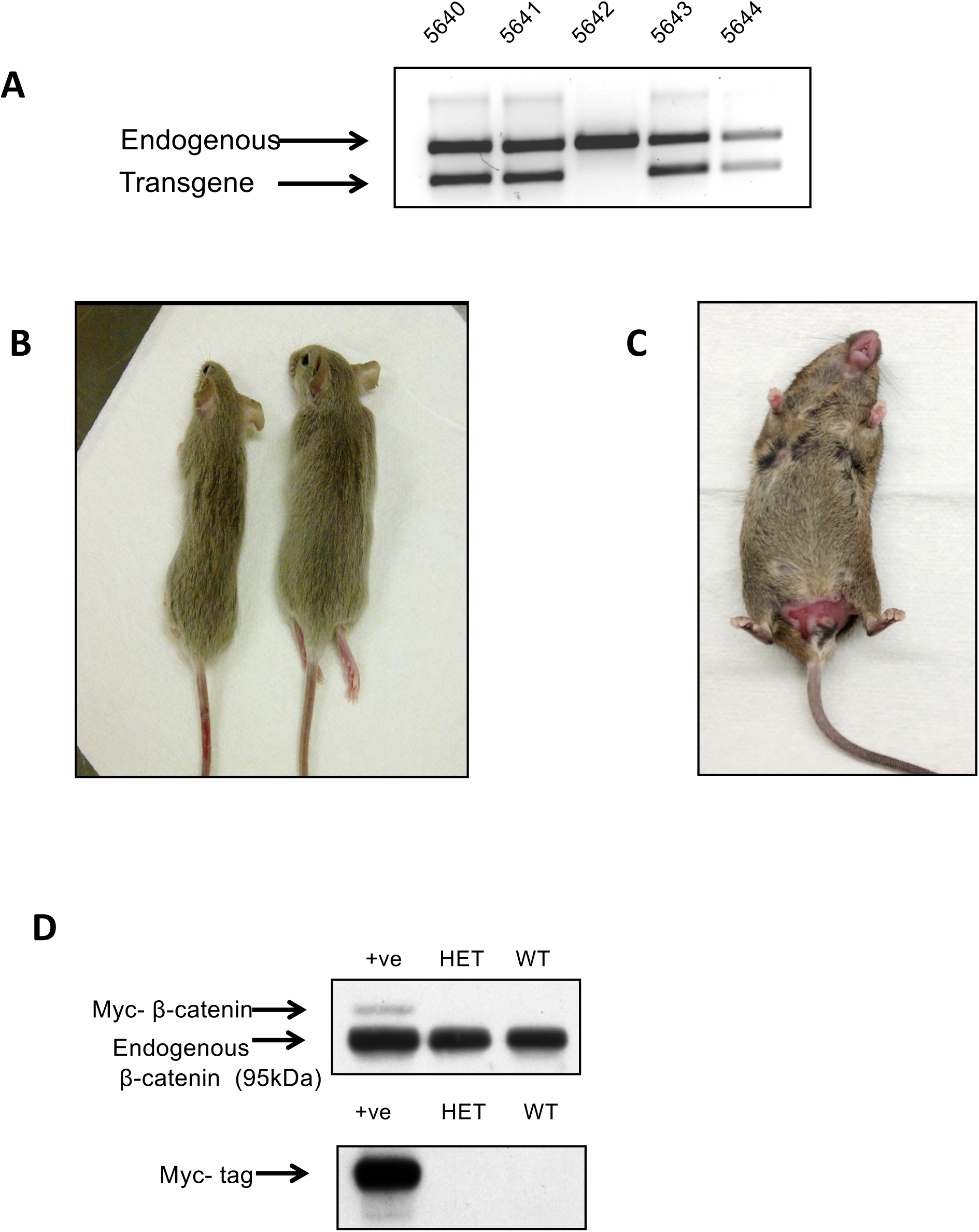
Assessment of mice carrying bcatFL transgene. A: genotyping of F1 generation pups for presence of bcatFL transgene. B: comparison of size of mice carrying 1 allele of bcatFL (left) to wildtype littermates (right). C: image of skin condition that developed in mice carrying the transgene. D: immunoblot of liver lysates from mice carrying 1 allele of bcatFL transgene that were negative for Cre expression.

Full necropsies performed on these mice failed to uncover a specific mediator of the condition; however, placing breeding pairs on doxycycline chow slowed the development of the erosion, allowing healthy litters to be born. These mice were of equal size to their wildtype littermates and exhibited no evidence of the skin condition. It remains undetermined as to what causes the skin erosion to develop. We have evidence that it is not transgene specific as 2 collaborating labs working on similar projects and using the same plasmid system also had mice that developed this condition. We were able to move forward with our study by placing all breeding cages on doxycycline chow, with pups placed on normal chow at weaning. Following this procedure, transgene positive mice were healthy, but the condition would recur if the breeding pairs were taken off doxycycline containing chow.

### Is stabilized β-catenin required for long-term maintenance of mammary tumor growth?

To assess the role of stabilized β-catenin in the maintenance of tumor growth following tumor initiation, 6 animal cohorts were established, along with control animals. Two different mammary Cre strains, WAP-Cre and MMTV-NLST-Cre, were crossed with the regulatable, stabilized β-catenin (bcatFL) animals, as well as with animals expressing mutated p53-R270H. These cohorts were designed to assess whether the importance of stabilized β-catenin to continued tumor growth was altered when mice contained a second driving mutation. However, the growth kinetics and penetrance were far lower than anticipated or has been seen with other models of stabilized β-catenin expression, with or without altered p53 expression. In the MMTV cohort, very few animals expressing stabilized β-catenin developed tumors, and those tumors that did arise required almost 18 months (Figure 3A). In comparison, stabilized β-catenin expression targeted to WAP-Cre expressing mammary epithelial cells resulted in a greater penetrance than with MMTV. Still, only 24% of mice generated expressing bcatFL and WAP-Cre developed tumors, on average 310 days after the birth of their first litter (Figure 3B). The double transgenic mice, expressing both stabilized β-catenin and mutated p53, targeted with either MMTV or WAP-Cre showed similar survival curves to that of mice expressing p53 alone, with no co-operation observed between the 2 tumor drivers (Figure 3A and B).

**Figure 3.**
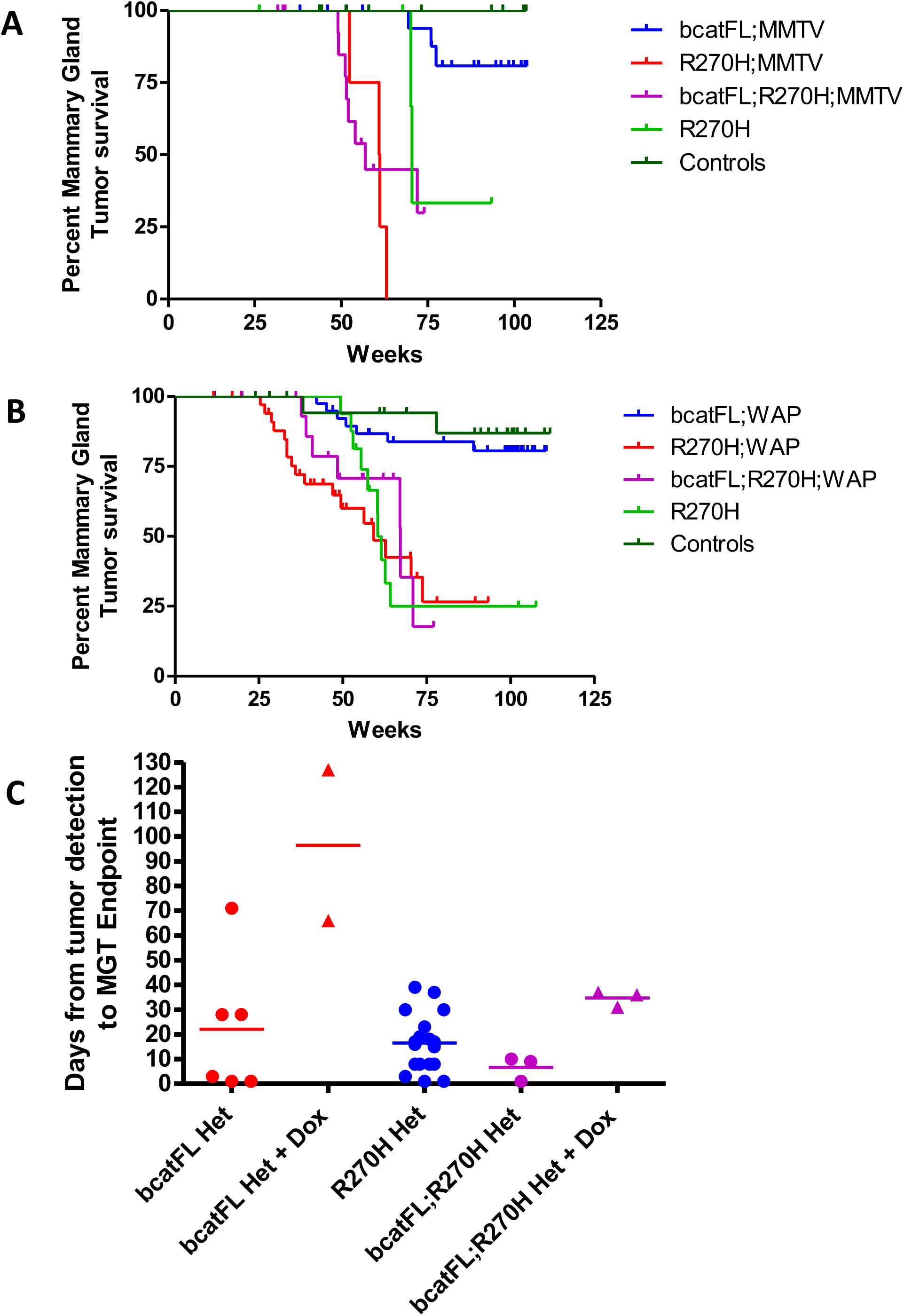
Survival analysis. A: Kaplan-Meier (K-M) plot of animal cohorts generated with MMTV-NLST-Cre (bcatFL;Cre, n=20, R270H;Cre, n=4, bcatFL;R270H;Cre, n=17, R270H+/−, n=5, and Controls, n=16). B: K-M plot of animal cohorts generated with WAP-Cre (bcatFL;Cre, n=40, R270H;Cre, n=36, bcatFL;R270H;Cre, n=16, R270H+/−, n=16, and controls, n=21). C: impact of turning off bcatFL transgene expression on the rate of tumor growth following tumor initiation in WAP-Cre driven tumors (bcatFL;Cre, n=6, bcatFL;Cre + Dox, n=2, R270H;Cre, n=18, bcatFL;R270H;Cre, n=3, and bcatFL;R270H;Cre + Dox, n=3).

Although the number of mice that developed mammary tumors was significantly lower than expected, we explored the importance of stabilized β-catenin expression to long-term maintenance of mammary tumor growth where possible. As seen in Figure 3C, when 2 animals with bcatFL tumors were placed on doxycycline we observed that tumor growth slowed upon extinguishing bcatFL expression. This is similar to what was observed when the bcatFL/p53 mice were placed on doxycycline. In both cases tumor stasis was observed for a period before tumor growth resumed and the mice eventually reached tumor size endpoint. None of the mice showed regression of the tumor upon silencing of bcatFL expression.

While no cooperation was observed between stabilized β-catenin and p53 in our mouse model, we examined the tumor histology to determine if the pattern of histological subtypes differed between the single and double transgenic mice. The histology of all 3 groups, bcatFL, p53-R270H and bcatFL; R270H generated with WAP-Cre, were predominately of the adenosquamous carcinoma type, with tumors filled with keratin. One difference observed between the p53-R270H alone and bcatFL; R270H tumors is that none of the tumors from the bcatFL; R270H mice were classified as poorly differentiated adenocarcinomas, which is in stark difference to the 30% of p53-R270H tumors that are classified in this group (Figure 4).

**Figure 4.**
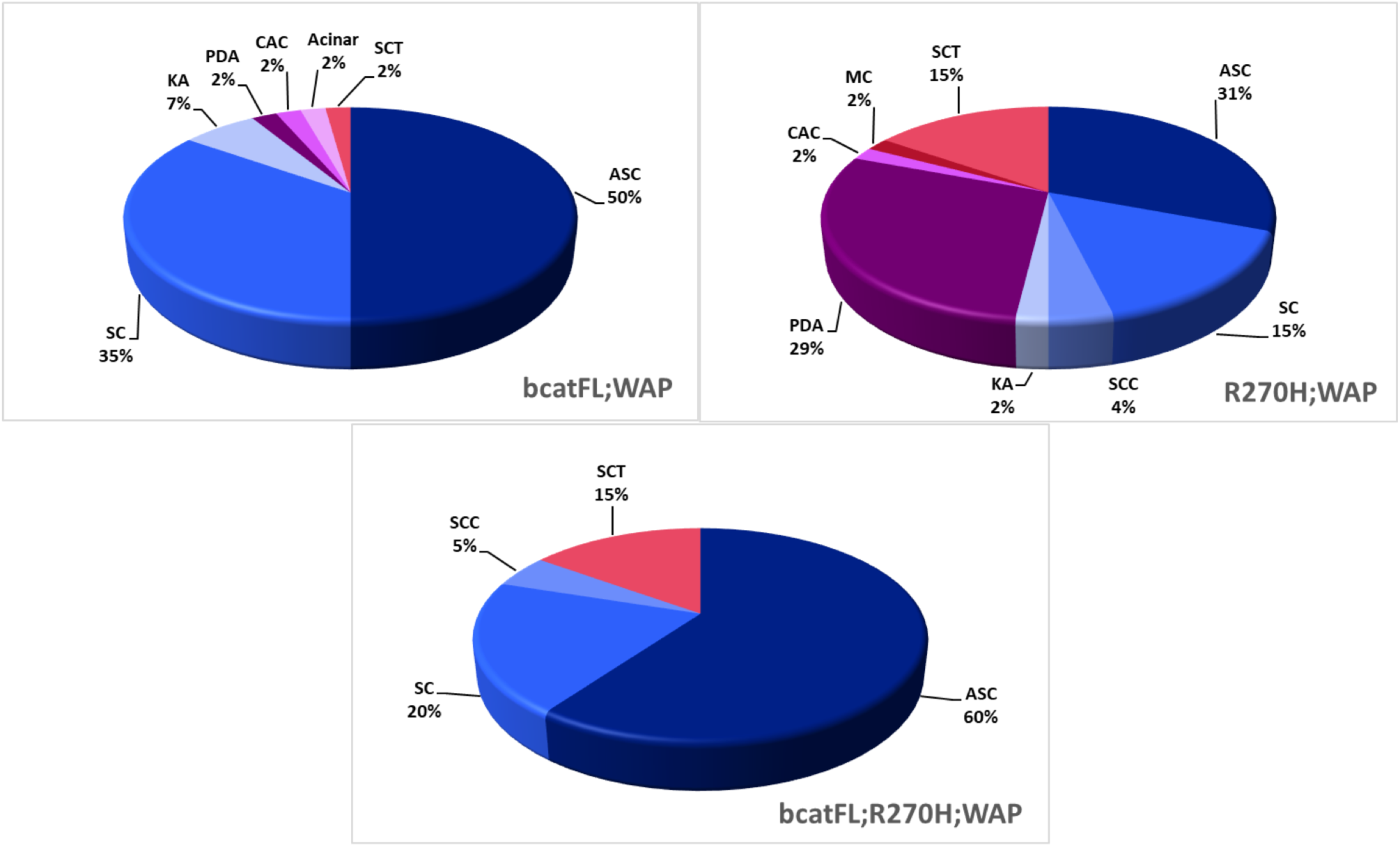
Histological assessment of WAP-Cre driven tumors. ASC=adenosquamous carcinoma, SC=squamous cyst, KA=keratoacanthoma, PDA=poorly differentiated adenocarcinoma, CAC=complex adenocarcinoma, Acinar=acinar adenocarcinoma, SCT=spindle cell tumor, SCC=squamous cell carcinoma, and MC=metaplastic carcinoma. bcatFL;Cre, n=46 tumors, R270H;Cre, n=52 tumors and bcatFL;R270H;Cre, n=20 tumors.

## Discussion

At the outset of this project, a mouse model of regulatable activated β-catenin had not been established. Since that time, a doxycycline-on regulatable ΔN89-β-catenin mouse model was reported, with the addition of luciferase co-expression (17, 21). These authors observed development of β-catenin-driven mammary tumors, the majority of which regressed upon doxycycline removal. The 10% of tumors that failed to regress did not express the ΔN89-β-catenin and therefore were presumably driven by a secondary, undetermined mutation (21). The study did not assess whether growth was re-initiated in regressed tumors. The poor penetrance of tumors observed in the model presented here may be a consequence of the need to maintain breeding pairs on doxycycline until weaning to avoid the skin phenotype as that also likely suppresses initial expression of β-catenin once the Cre drivers are activated. This might suggest that there is a window of tumor initiation susceptibility very early in mammary gland development. We do not know why the strategy used here led to the skin erosion phenotype but this was not β-catenin dependent as it also occurred with other gene cargoes. Since it was suppressible by doxycycline, we surmise the effect is related to the integrated pTET-BigT plasmid sequences.

In summary, our data are consistent with β-catenin playing a role in tumor initiation but given the small number of tumor-bearing animals resulting from the study, we cannot assess its role in long-term maintenance of mammary tumors.

## Materials and Methods

### Targeting Vector Construction

Stabilized β-catenin cDNA in the pcDNA3 plasmid was previously generated in the Woodgett lab with amino acids S33, S37, T41 and S45 mutated to alanine. A partial fragment of this stabilized β-catenin cDNA (up to XbaI cut site) was excised and inserted in frame into a pcDNA3 vector containing fireflluciferase cDNA followed by a T2A sequence using the Infusion HD-eco dry kit (Clontech), as per the manufacturer’s instructions. This Luciferase-T2A-partial β-catenin sequence was then excised and ligated into the NheI/XhoI fragment of the pTet-BigT vector (a gift of Dr. Andy McMahon). The remaining portion of the stabilized β-catenin cDNA was then ligated into the pTet-BigT vector following digestion with NdeI and XhoI. The completed pTet-BigT-luciferase-β-catenin sequence was then excised with PacI and AscI and ligated into the ROSA26 PAM1 vector, which had also been digested with PacI and AscI. The targeting vector was then linearized with SwaI to prepare for ES Cell electroporation.

### ES Cell Culture

R1 mouse embryonic stem cells were cultured in DMEM containing 15% FBS, 1mM Sodium Pyruvate, 100uM non-essential amino acids, glutamine, pen/strep, β-mercaptoethanol and LIF on mitomycin C treated mouse embryonic fibroblasts (MEF). Electroporation of the bcatFL targeting vector was carried out on R1 cells at passage 13 or 14 and G418 (160ug/mL) selection media was added approximately 24 hours after electroporation. Resistant colonies were then transferred to feeder MEF coated 96 well plates to grow out in order to screen for homologous recombination.

### Screening for Homologous Recombination

G418-resistant ES cell clones were screened via PCR using a forward primer (5’-CGC CTA AAG AAG AGG CTG TG-3’) located in the endogenous Rosa26 locus (not found in the targeting vector) and a reverse primer (5’-GAA AGA CCG CGA AGA GTT TG-3’) located within the tet machinery of the targeting vector. PCR was carried out using the Qiagen hot start Taq kit according to manufacturer’s instructions. Briefly, after 15 min initial denaturation at 95°C, 40 cycles of 94°C for 1 min followed by annealing at 49°C 1 min and elongation at 72°C for 2 mins was performed for the first cycle with an step increase in annealing temperature of an additional 3°C at cycle 2, 11 and 21, to yield a 1.3 kB band.

### Testing for proper transgene expression

The H5 and H22 ES cell clones were transfected with the pCAGGS-Cre plasmid to remove the lox-stop-lox preventing firefly luciferase and stabilized β-catenin expression. Puromycin resistant clones were collected and grown out to test for expression of both transgenes by immunoblotting.

### Southern Blotting

DNA was extracted from ES cell clones A2, H5 and H22, which were positive for homologous recombination by PCR and an untargeted control clone. Following extraction, 20ug of DNA was digested with EcoRV and ClaI to distinguish between the endogenous Rosa26 sequence, which yields a 11kb band and the inserted targeting vector containing the ClaI site and yielding a 9.7kB band. As a positive control, the pROSA 3’ probe plasmid was digested with XhoI to yield a 4kB linear fragment and 3 separate samples were run with 2, 10 or 25pg of DNA. The Southern blot was performed as described in Adams *et al.* (2).

### Chimera Generation, Animal Husbandry & Genotyping

2 separate clones, H5 and H22, were selected to undergo diploid aggregation in ICR mice. Chimeras of differing percentages were then crossed with FVB and C57Bl/6 mice to assess germline transmission. Tail or ear clips were digested in Proteinase K containing digest buffer (1x PCR buffer (ThermoFisher), 0.45% NP-40, 0.45% Tween20, and 1mg/mL Proteinase K (Invitrogen)) for 3.5 hr at 60°C, followed by 15 minutes at 95°C. Genotyping PCRs were performed using 10x buffer (B33, ThermoFisher), 10mM dNTP (Frogge Bio), 10µM primers (200nM final) (IDT Tech), and Taq polymerase (made in house). Animals were genotyped for the presence of all transgenes using the primers listed below.

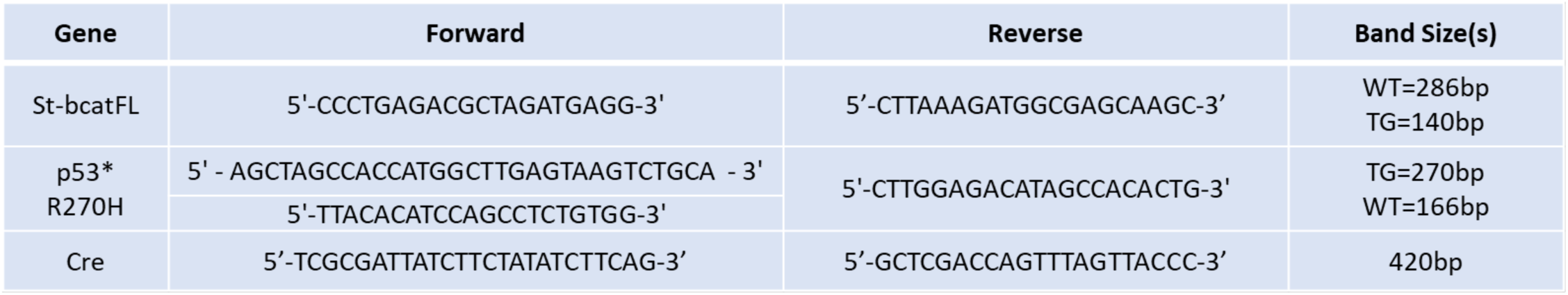

### Western Blot

Cell or tissue extracts were run out on an SDS PAGE gel, followed by transfer to a PVDF membrane. Blots were blocked in 5% skim-milk TBST for 1 hr prior to primary antibody incubation overnight at 4°C. including β-catenin (1:5000, BD Biosciences), Luciferase (1:2500, Promega), Myc tag (1:1000, Santa Cruz), and GAPDH (1:250,000, AbCam). Following overnight primary antibody incubation and TBST washes, blots were incubated with either anti-goat (1:5000, BioRad), rabbit (1:5000, GE) or mouse (1:5000, GE) secondary antibodies diluted in 5% skim-milk TBST for 1 hr, followed by both TBST and TBS washes. Blots were incubated with chemiluminescent reagents (Millipore) and exposed to film (GE).

### Pathology

Full necropsy on animals carrying one allele of the bcatFL transgene were carried out by pathologists at The Centre for Phenogenomics (www.phenogenomics.ca). Tissue embedding, sectioning and H & E staining were carried out by the Histology Core at The Centre for Phenogenomics.

## Acknowledgements

The authors would like to acknowledge Gessica Rapioni, Amanda Pagala and Kelly Jackson from The Centre for Phenogenomics for their assistance with monitoring animals for the development of the skin erosion condition. They also would like to acknowledge the Pathology Core at The Centre for Phenogenomics for their technical services.

